# Inhibition of PTCH1 drug efflux activity enhances chemotherapy efficacy against triple negative breast cancer

**DOI:** 10.1101/2025.09.03.673759

**Authors:** Sarah Cogoluegnes, Sandra Kovachka, Thierry Dubois, Roberto Würth, Elisa Donato, Andreas Trumpp, Michel Franco, Frédéric Luton, Stéphane Azoulay, Isabelle Mus-Veteau

## Abstract

Triple-negative breast cancer (TNBC) is the most aggressive breast cancer subgroup characterized by a high risk of resistance to chemotherapies and high relapse potential. High levels of mRNA from the Hedgehog receptor PTCH1 are associated with poor prognosis in TNBC. PTCH1 is overexpressed in many aggressive cancers. We previously reported that PTCH1 is a multidrug transporter that triggers resistance to chemotherapy of adrenocortical carcinoma and melanoma cells, and that inhibiting PTCH1 drug efflux strongly enhanced chemotherapy efficacy on these cell lines both *in vitro* and *in vivo*. In the present study, we found that PTCH1 inhibition also significantly inhibited doxorubicin efflux in three TNBC cell lines leading to a strong increase of the cytotoxic effect of doxorubicin and docetaxel, and an inhibition of cell migration. Altogether, our data highlight the therapeutic potential of targeting PTCH1 drug efflux activity using drug association strategies for the treatment of TNBC patients.

## Introduction

Breast cancer (BC) is the most prevalent tumor and the main cause of cancer death in women worldwide with approximately 2.2 million new cases and 685 000 deaths reported globally in 2020 (Sung et al., 2021, Arnold et al., 2022). It encompasses a complex spectrum of diseases with different genetic and molecular features contributing to tumor development and progression (Shiovitz and Korde, 2015, Gradishar, 2023). Invasive breast cancer (IBC) have been classified using histological evaluation, molecular profiling and surrogate markers. The World Health Organization (WHO) recognizes 18 distinct histological subtypes of IBC, with invasive carcinoma of no special type (NST) being the most common, accounting for 40% to 80% of cases (Łukasiewicz et al. 2021). IBC can be subdivided into molecular subtypes, namely human epidermal growth factor receptor 2 (HER2) enriched luminal A, luminal B, and basal-like, based on tumor gene-expression patterns. In clinical practice, surrogate markers such as estrogen receptor (ER), progesterone receptor (PR), and HER2, assessed through immunohistochemistry (IHC), are commonly used for tumor classification (Carey et al 2006). 15–20% of all breast cancer cases lacking ER and PR expression and *HER2* gene amplification, are classified as triple-negative breast cancer (TNBC). These subtypes exhibit distinct biological characteristics, clinical outcomes, and treatment responses (Johnson et al 2021). Many approaches have been employed for breast cancer treatment. Surgery is an important way for breast cancer treatment, with options including breast-conserving surgery or mastectomy (Christiansen et al 2022). Chemotherapy may be administered either before (neoadjuvant) or after (adjuvant) surgery and is tailored to the tumor’s specific characteristics (Rouzier et al 2005). Radiation therapy destroys the remaining cancer cells after surgical treatment or chemotherapy, thereby improving overall survival (OS) rates and decreasing recurrence (Luz et al 2022). Endocrine therapies target ERs by either reducing estrogen levels or blocking estrogen-induced stimulation of breast cancer cells (Patel et al 2023). However, approximately 50% of hormone receptor-positive breast cancers are resistant to this treatment (Drãgãnescu and Carmocan 2017). Due to the absence of relevant receptor markers, patients with TNBC do not benefit from established endocrine or HER2-targeted therapies, and general chemotherapy remains the standard treatment for nonsurgical TNBC. However, less than 30% of patients achieve a complete response, and both recurrence and mortality rates remain elevated compared to non-TNBC subtypes (Li Y et al 2022). Although BC survival has improved over the last decades, treatments resistance still represents a significant unmet clinical need (Gradishar 2023).

Analogous to other developmental pathways, aberrant Hedgehog (HH) pathway activity is critical for the initiation and progression of various tumors types (Scales and de Sauvage 2009). O’Toole and coworkers (2011) observed that the expression of Hedgehog ligand such as SHH was significantly associated with increased risk of metastasis, breast cancer-specific death, and a basal-like phenotype in a cohort of patients with invasive ductal carcinoma of the breast. Their data suggested that epithelial–stromal HH signaling, driven by ligand expression in carcinoma cells, promoted breast cancer growth and metastasis. Jeng and coworkers (2013) also observed on a cohort of patients with invasive breast carcinoma following curative resection that, compared with paired noncancerous tissue, a higher expression of mRNAs from the HH pathway components SHH, PTCH1, GLI-1, and SMO in breast cancer tissue correlates with invasiveness and is associated with an increased risk of recurrence.

We discovered that the receptor of SHH, PTCH1, is a multidrug transporter able to efflux chemotherapeutic agents out of cancer cells, which contributes to chemotherapy resistance of adrenocortical carcinoma and melanoma cells (Bidet et al 2012, Hasanovic et al 2018, Signetti et al 2020, Feliz Morel et al 2022). We showed that PTCH1 uses the proton-motive force to efflux drugs like the bacterial efflux pumps from the RND family (Bidet et al 2012). Therefore, PTCH1 is active as a drug efflux pump only in cancer cells which exhibit an acidic extracellular pH due to their high glucose consumption in contrary to normal cells which exhibit an extracellular pH slightly more basic than intracellular one (Damaghi et al 2013). Moreover, in adults, PTCH1 is expressed very slightly. This makes PTCH1 a highly relevant therapeutic target which inhibition should affect only cancer cells in contrary to the multidrug transporters from the ABC family such as the P-glycoprotein (P-gp) which inhibition is toxic for healthy cells (Kathawala et al. 2015). The screening of about 2000 small molecules led us to identify a the panicein A hydroquinone (PAH) extracted from the marine sponge *Haliclona mucosa* as an inhibitor of PTCH1 drug efflux activity (Fiorini et al 2015). We developed the synthesis of PAH and showed that the synthesized PAH increased the efficacy of conventional chemotherapies like doxorubicin and that of targeted therapies such as vemurafenib, an inhibitor of the BRAF mutant V600E, on BRAFV600E-expressing melanoma cells both *in vitro* and *in vivo* (Signetti et al 2020, Kovachka et al 2021, Kovachka et al 2022).

In the present study, we reported the mRNA expression levels of PTCH1 in several cohorts of breast cancer patients, in isolated circulating tumor cells from blood samples from patients with metastatic breast cancer, and in several types of breast cancer cell lines. We observed that PTCH1 is more strongly expressed in TNBC. We found that PTCH1 inhibition significantly inhibited chemotherapy efflux in three TNBC cell lines leading to a strong increase of the doxorubicin and docetaxel efficacy. Our data highlight the therapeutic potential of targeting PTCH1 drug efflux activity to enhance the treatment efficacy for TNBC patients.

## Materials and methods

### Chemical and biological material

Panicein A hydroquinone was synthesized as described in (Signetti et al 2020).

Doxorubicin hydrochloride and docetaxel were purchased from Sigma-Aldrich and Acros-Organics, respectively.

Human breast cancer cell lines MDA-MB-231, MDA-MB-468 and HCC-38 were purchased from ATCC. Cells were cultured at 37°C in a 5% CO_2_ /95% air water-saturated atmosphere in DMEM medium supplemented with 10% fetal bovine serum and penicillin/streptomycin for MDA-MB-231, in DMEM-F12 medium supplemented with 2 mM HEPES, 10% fetal bovine serum and penicillin/streptomycin for MDA-MB-468, and in RPMI medium supplemented with 1.5 g/L sodium bicarbonate, 10 mM HEPES, 1 mM sodium pyruvate, 10% fetal bovine serum and penicillin/streptomycin for HCC-38.

K699 *S. cerevisiae* yeast strain (Mata, ura3, and leu 2–3) and K699 S. c. strain expressing human PTCH1 were grown as described (Joubert et al 2009).

### PTCH1 mRNA expression analyses in human biopsies and in human breast cancer cell lines

TCGA cohort: the publicly available RNA-SeqV2 Level 3 dataset (January 2015) were downloaded from The Cancer Genome Atlas (TCGA) breast invasive carcinoma cohort (http://cancergenome.nih.gov/) (Cancer Genome Atlas Network 2012) and integrated into a platform in knowledge data integration (KDI) at Institut Curie (https://bioinfo-portal.curie.fr). We classified the breast cancer subgroups, based on the immunohistochemical status for ER, PR and HER2, as described (Suresh et al 2022). TNBC (ER-, PR-, HER2-), HER2 (ER-, PR-, HER2+), luminal A (ER+ and/or PR+, HER2-), and luminal B (ER+ and/or PR+, HER2+). Cell lines: total RNA from the different breast cancer cell lines was extracted and hybridized to GeneChip™ Human Exon 1.0 ST arrays (Affymetrix) as previously described (Dakroub et al 2023).

Using breast cancer gene-expression miner (bc-GenExMiner v4.5): this is a breast cancer-associated web portal (http://bcgenex.ico.unicancer.fr) with a statistical mining module allowing several differential gene expression analyses based on microarray or RNAseq transcriptomic data on sixty-two breast cancer cohorts (Jezequel et al 2021). Prognostic analyses were performed on this web portal using exhaustive prognostic analysis which permits to screen the prognostic impact of a gene or a specific Affymetrix® probeset ID on all possible combinations of population (significant results may be considered robust if more than 5 combinations among the 27 give a significant result. If there are only 5 combinations or less with a p-value<0.05, one cannot exclude a false discovery problem) (Fig. 1), and intrinsic molecular subtypes prognostic analysis which permits to assess the prognostic impact of a gene or a specific Affymetrix® probeset ID within groups of patients with a certain intrinsic molecular subtypes as defined by Single Sample Predictors (SSP) and/or Subtype Clustering Models (SCM) (Sup Fig 1).

**Figure 1.**
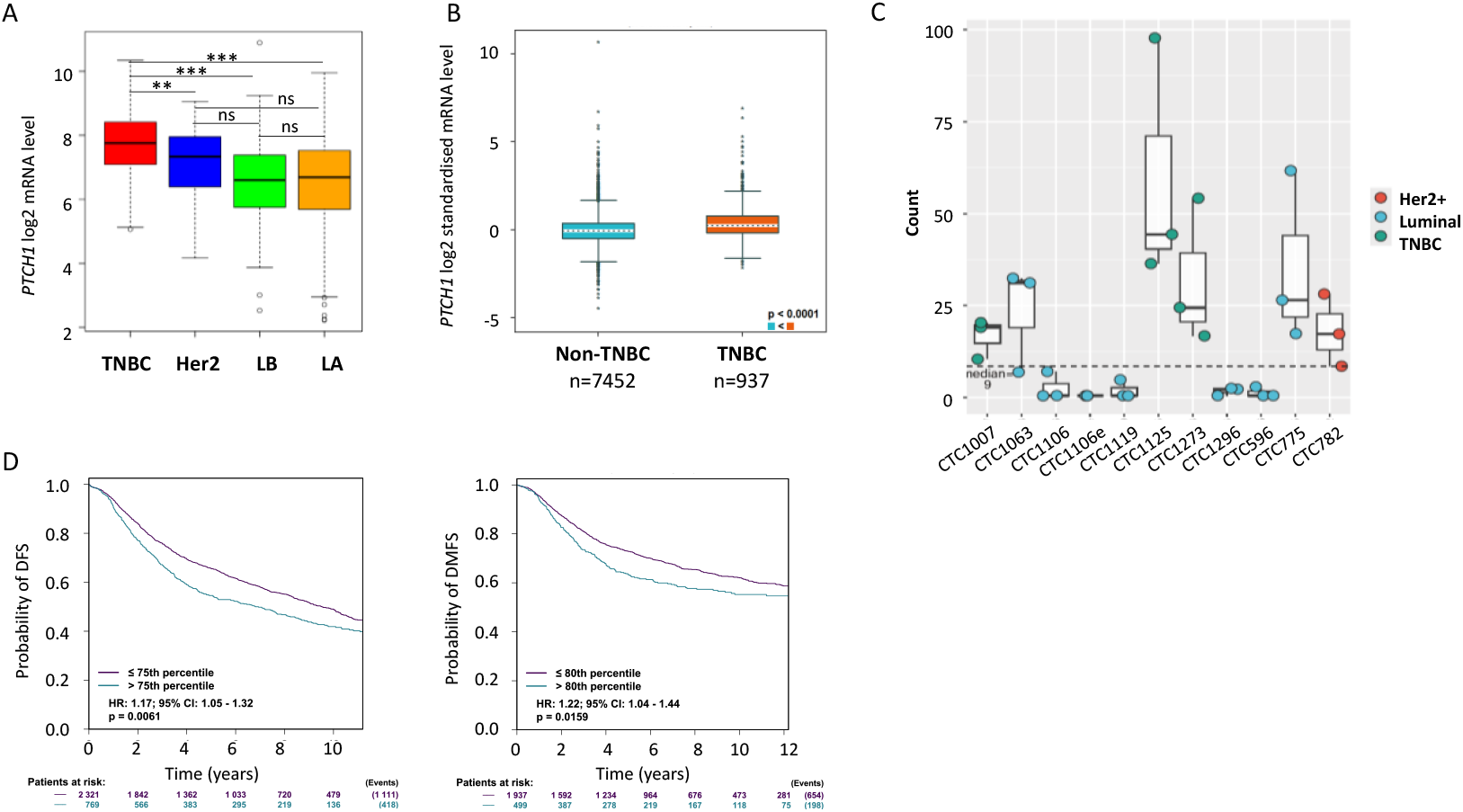
PTCH1 is expressed in breast cancer patients. **A**. Relative *PTCH1* RNA expression in TNBC (red), Her2 (blue), luminal B (LB, green), and luminal A (LA, orange) in the TCGA cohort is illustrated by box plots (log_2_ transformed). Outliers are shown within each population (open circles). Student’s t test was used to compare RNA levels between two groups. The p values are indicated (*P < 0.05; **P < 0.01; ***P < 0.001; ns P > 0.05). **B**. Normalized *PTCH1* mRNA expression according to TNBC (IHC) status from all DNA microarray data from bc-GenExMiner v4.5 illustrated by box plots (log_2_ transformed) (Jezequel et al 2021). **C**. Boxplots showing *PTCH1* mRNA expression (normalized counts from RNA seq data) in circulating tumor cells (CTC) isolated from metastasis breast cancer patient peripheral blood (Wurth R et al 2025). **D**. High *PTCH1* mRNA expression is associated with a poorer prognosis in breast cancer. Distant Metastasis Free Survival (DMFS) and disease-free survival (DFS) data based on *PTCH1* mRNA expression were obtained using the exhaustive prognostic analysis on all status breast cancers on the bc-GenExMiner v4.5 web portal, and illustrated by Kaplan–Meier (KM) curves for all ER and PR breast cancers with positive nodules. The obtained Hazard Ratio (HR) with 95% confidence interval and log-rank p-values are shown.

### PTCH1 expression analysis on organoids derived from circulating tumor cells from breast cancer patients

For the analysis of *PTCH1* expression on CTC-derived organoids from metastatic breast cancer individuals, data from Würth and colleagues (2025) were re-analyzed with DESeq2 package (v1.26.0) using RStudio (v1.4).

### SDS-PAGE and western blotting

Total RIPA extracts from cells were prepared. Protein concentrations were determined by the Bradford Protein Assay (Bio-Rad). Samples (50 to 80 µg) were separated on SDS-PAGE and transferred to nitrocellulose membranes (Amersham) using standard techniques. After 1 h at room temperature in blocking buffer (20 mmol/L Tris-HCl pH 7.5, 45 mmol/L NaCl, 0.1 % Tween-20, and 5 % non-fat milk), nitrocellulose membranes were incubated overnight at 4 °C with the monoclonal rat anti-PTCH1 antibody from R&D system biotechne clone 413220 (1/1000), the monoclonal mouse anti-P-gp antibody from abcam (ab3366, 1/1000), the rabbit anti-ABCG2 antibody from GeneTex (GTX50793, 1/500) or the rabbit anti-GAPDH antibody from Elabscience (1/20000). After 3 washes, membranes were incubated for 45 min with anti-rabbit (1:2000) or anti-mouse (1:5000) immunoglobulin coupled to horseradish peroxidase (Dako). Detection was carried out with an ECL Prime Western Blotting detection reagent (SuperSignal West Femto Maximum Sensitivity Substrate from ThermoScientific) on a Fusion FX imager (Vilber Lourmat), and analyses were performed using ImageJ software.

### Cytotoxicity assays

#### On adherent cells

Cells were seeded in 96-well plates in triplicate and grown in medium to achieve 70 % to 80 % confluence. Medium was then removed and replaced with 100 µL/well of complete medium containing PAH or DMSO as a control. After 15 min, 100 µL of complete medium containing serial dilutions of doxorubicin or docetaxel were added. Plates were incubated at 37 °C and 5 % CO_2_. After 24 or 48 hours, cells were incubated for 3 hours at 37 °C with 100 µL/well neutral red (NR) solution (50 µg/mL in medium) following the manufacturer’s protocol. Measurements were made in microplate readers (Multiskan Go Microplate Spectrophotometer from Thermo Scientific). IC_50_ was defined as the concentration that resulted in a 50 % decrease in the number of viable cells, and IC_50_ values were calculated using GraphPad Prism 6 software.

#### On spheroids

MDA-MB-231 cells were plated in ultralow attachment 24 well plates in complete medium containing PAH or DMSO as a control and 100, 200, 250 or 300 µM docetaxel. Plates were incubated at 37 °C and 5 % CO_2_. After two weeks, pictures of each well were taken using the Cytation 5 cell imaging system from Biotek.

#### Transwell invasion assay

The cell invasion assay was performed according to the manufacturer’s instructions (Costar) with a 24-well transwell plate containing transwell cell culture chamber inserts with polycarbonate membranes to study cell invasion. Briefly, 3 × 10^5^ cells per well were plated in the upper chamber of the transwell plate in complete culture medium containing 15 µM PAH or DMSO, and 0 or 1 µM doxorubicin in one plate and 0 or 20 µM docetaxel in the other one. Complete culture medium was added to the lower chamber, and transwell plates were incubated at 37°C in 5% CO_2_ atmosphere 24 hours for the one treated with doxorubicin or 48 hours for that treated with docetaxel. Then, the upper chambers were removed, and the wells (lower chambers) were fixed, stained with crystal violet and observed on a microscope with X5 objective.

#### Wound-healing assay

Once cells were confluent in 24-well plates, a wound was performed with a P200 pipette tip, and the medium was replaced by fresh medium containing or not 5 µM docetaxel or/and 5 µM PAH. Pictures of each well were taken immediately after wounding, and 72 hours after wounding with a Leica DM IRB (5X objective). The width of the wound was measured using ImageJ software and reported as final wound width /initial wound width in percentage.

#### Doxorubicin efflux measurements

Cells were seeded on coverslips in 24-well plates and allowed to grow to 80% confluence. Coverslips were incubated at 37 °C and 5 % CO_2_ with 10 μM doxorubicin in physiological buffer (140 mM NaCl, 5 mM KCl, 1 mM CaCl_2_, 1 mM MgSO_4_, 5 mM glucose, 20 mM HEPES, pH 7.4). After 2 hours, three coverslips were immediately fixed with 4 % PFA for the doxorubicin loading control, rapidly washed with PBS and mounted in SlowFade Gold antifade reagent with DAPI (Invitrogen). The other coverslips (triplicate per condition) were incubated with physiological buffer supplemented with DMSO or 10 µM of PAH under gentle shaking at room temperature and protected from light. After 30 min, coverslips were fixed with 4 % PFA, washed and mounted as described above. For competition on doxorubicin loading, MDA-MB-231 cells seeded on coverslips were incubated for 1 hour at 37 °C and 5 % CO_2_ with 10 μM doxorubicin in physiological buffer in the presence or the absence of 50 µM docetaxel. Images were acquired with a Zeiss Axioplan 2 fluorescence microscope coupled to a digital charge-coupled device camera using a 40X/1.3 Plan NeoFluar objective and filters for Alexa 594. Doxorubicin fluorescence was quantified using ImageJ software. Sampling of cells was performed randomly. About 100 cells (from three wells) were scored per condition per experiment.

#### Yeast growth inhibition by drugs

*S. cerevisiae* expressing wild-type human PTCH1 were pre-cultured minimum medium to OD_600nm_ = 2, diluted to OD_600nm_=0.2 in rich medium supplemented or not with 5 µM PAH and with 15 µM doxorubicin or 400 µM of docetaxel, and grown at 18°C in 96-well plates (BD Biosciences). Absorbance at 600 nm was recorded during growth.

#### Statistical analysis

All results represent at least three independent replications. Data are shown as mean value ± SEM. Prism 6 (GraphPad) was used to determine IC_50_ values and other statistical analyses using one-way analysis of variance (ANOVA) followed by Bonferroni’s Multiple Comparison Tests.

## Results

### PTCH1 mRNA is expressed in human breast cancer and in circulating tumor cells

Analysis of the cancer genome atlas (TCGA) cohort showed that *PTCH1* mRNA is expressed in all breast cancer subgroups, with the highest expression in TNBC (Fig. 1A). PTCH1 expression analysis using breast cancer gene-expression miner (bc-GenExMiner v4.5) (Jezequel et al 2021) also showed that *PTCH1* mRNA is significantly more expressed in TNBC compared to other breast cancers (Fig. 1B).

Furthermore, we analyzed *PTCH1* expression in circulating tumor cells (CTCs) by leveraging a recent dataset of RNA-seq analysis of CTC-derived organoids (CDOs) isolated from 11 liquid biopsies from 10 metastatic breast cancer patients, encompassing all the different major subtypes of breast cancer (Würth et al. 2025). *PTCH1* expression varied among CDOs, with null/low expression in 5 CDOs and higher expression in 6 CDOs. Notably, higher *PTCH1* expression was observed in 100% (3/3) of CDOs established from TNBC patients (Fig. 1C). These data indicate that *PTCH1* is expressed not only in patient breast tumors but also in patient CTC which are well known to drive metastasis, the leading cause of death in individuals with breast cancer.

We then examined whether *PTCH1* mRNA expression levels could have a clinical significance using prognostic analyses performed on breast cancer gene-expression miner web portal. The exhaustive prognostic analysis, which permits to screen the prognostic impact of PTCH1 on all possible combinations of population, revealed that high *PTCH1* mRNA expression was associated with lower distant metastasis free survival (DMFS) and disease-free survival (DFS) in 14 combinations among 27 and 7 combinations among 27 respectively, but not with lower overall survival (OS). This is illustrated in Figure 1D with Kaplan-Meier curves obtained on patients with all ER and PR status and node positive BC. However, the intrinsic molecular subtypes prognostic analysis on patients with high proliferative ER+/HER2-BC revealed that high *PTCH1* mRNA expression was associated with lower DMFS, DFS and OS (Sup Fig 1).

Transcriptomic analysis of 23 breast cancer cell lines revealed variable *PTCH1* mRNA levels depending on the cell line, with a tendency of highest levels in TNBC cell lines compared to other breast cancer cell lines (Fig. 2A). Western-blot analysis confirmed the expression of PTCH1 protein in 3 TNBC cell lines (MDA-MB-231, HCC-38 and MDA-MB-468), and showed that HCC-38 cells express significantly higher levels of PTCH1 protein compared to MDA-MB-231 and MDA-MB-468 cells (Fig. 2B).

**Figure 2.**
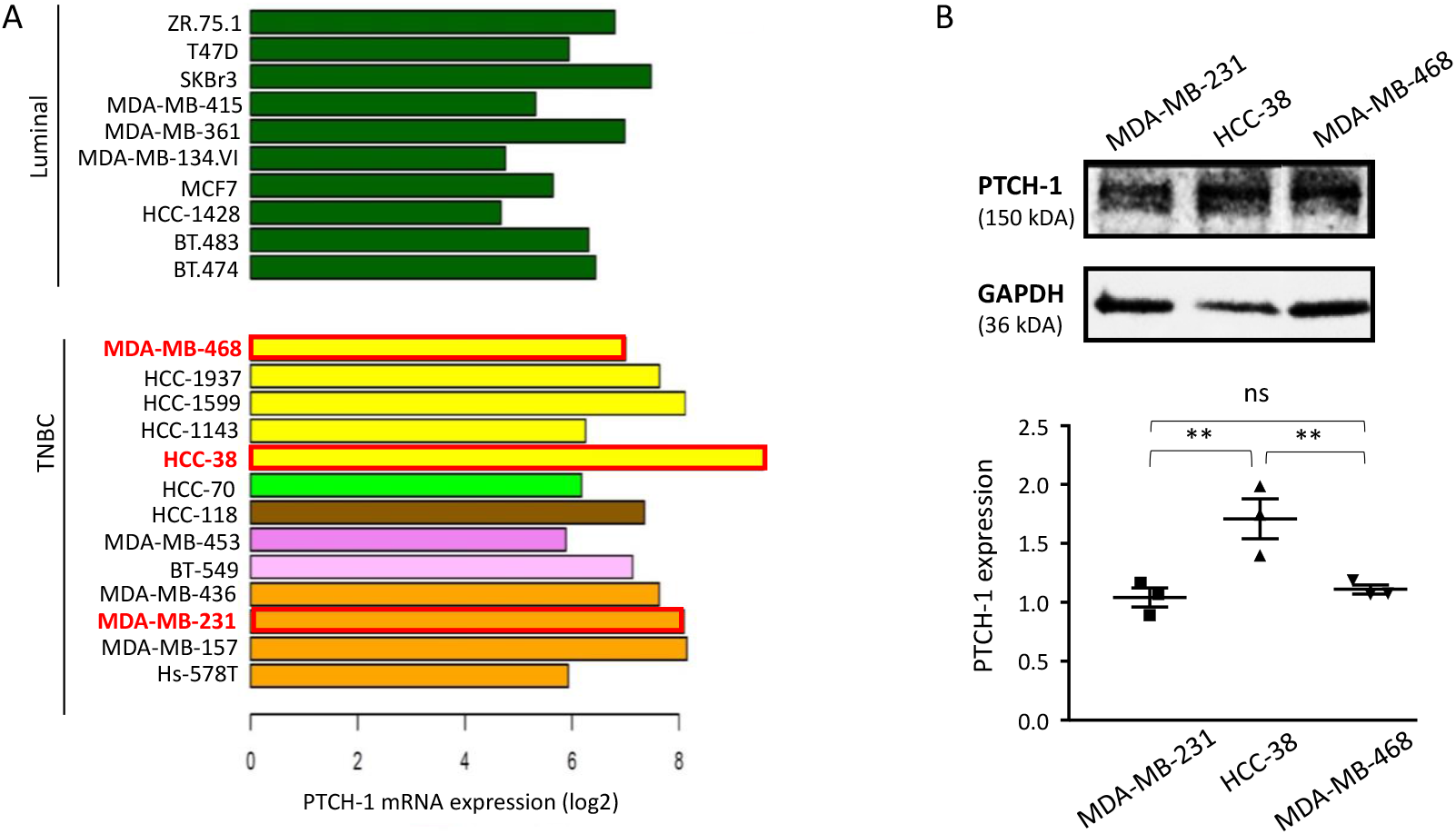
PTCH1 is expressed in breast cancer cell lines. **A**. *PTCH1* mRNA expression (log_2_ transformed) in various luminal and HER2 breast cancer cell lines (dark green) and in TNBC cell lines (other colors). TNBC cell lines are depicted according to the “Lehmann TNBC subtype” nomenclature (Lehmann et al 2011): basal-like 1 (yellow), basal-like 2 (pale green), immunomodulatory (brown), luminal androgen receptor (dark pink), mesenchymal (pale pink) and mesenchymal stem-like (pink). **B**. PTCH1 protein expression in three TNBC cell lines. Western-blot was performed on 50 µg extracts from TNBC cell lines (MDA-MB-231, HCC-38 and MDA-MB-468) with antibodies directed against PTCH1. PTCH1 and GAPDH signals were quantified using ImageJ software. Data presented are the mean ± SEM of at least 3 independent experiments. Significance is calculated using Oneway ANOVA Turkey’s multiple comparisons test and attained at P < 0.05 (*).

These studies confirm that PTCH1 is expressed in breast cancer and more particularly in TNBC. Therefore, if PTCH1 is involved in the resistance of TNBC to the conventional chemotherapies currently used to treat TNBC patients such as doxorubicin and docetaxel, it could be very interesting to inhibit it to increase the efficacy of these treatments.

We previously established a *Saccharomyces cerevisiae* yeast strain overexpressing human PTCH1, and we observed that these yeasts were able to grow in the presence of a doxorubicin concentration that prevents parental yeast strain to grow and to efflux more doxorubicin than parental yeast strain (Bidet et al 2012, Sup Fig. 2A). To know if PTCH1 is able to efflux docetaxel as well as doxorubicin, we cultivated the yeasts overexpressing PTCH1 in the presence of docetaxel. We show here that the expression of PTCH1 allows yeasts to better grow in the presence of docetaxel than parental yeasts (Sup Fig. 2B), which indicates that PTCH1 transports docetaxel out of yeasts. Moreover, the addition to the medium of the inhibitor of PTCH1 drug efflux activity (PAH) inhibits the growth of the yeasts expressing PTCH1 in the presence of docetaxel as it does in the presence of doxorubicin (sup Fig. 2). This observation suggests a role of PTCH1 in the efflux of docetaxel, and that targeting PTCH1 could be a good strategy to increase docetaxel efficacy in TNBC cells.

### Inhibition of PTCH1 drug efflux increases the sensitivity of TNBC cells to chemotherapies in 2D and in 3D culture

We then tested the effect of doxorubicin and docetaxel on the cell viability of three TNBC cell lines presenting an endogenous expression of PTCH1 (MDA-MB-231, HCC-38 and MDA-MB-468) as shown in Figure 2B. Cells were grown to 80% confluency, and treated with doxorubicin or docetaxel 24 or 48 hours respectively in the absence or the presence of the PTCH1 drug efflux inhibitor PAH. As reported in Figure 3, the addition of PAH to doxorubicin or docetaxel increased the cytotoxicity of both chemotherapies and significantly decreased their IC_50_ on the three cell lines in 2D culture (Table 1).

**Table 1.** PTCH1 drug efflux inhibitor PAH increases the cytotoxicity of docetaxel and doxorubicin against TNBC cells. Cell viability was measured after 24 hours or 48 hours treatment with increasing concentration of doxorubicin or docetaxel respectively on MDA-MB-231, MDA-MB-468 and HCC-38 cell lines in the absence or the presence of 15µM PAH. IC_50_ values of chemotherapy (corresponding to the concentration of chemotherapy inducing 50% of cell death) were calculated. Data reported are the mean ± SEM of at least 3 independent experiments.

|  |  | MDA-MB-231 |  | MDA-MB-468 |  | HCC-38 |  |
| --- | --- | --- | --- | --- | --- | --- | --- |
|  | PAH (μM) | 0 | 15 | 0 | 15 | 0 | 15 |
| Doxorubicin | IC<sub>50</sub> (μM) | 16.6±2 | 2.3±0.3 | 33.6±1.4 | 4.8±2 | 47.8±12 | 12±0.8 |
|  | Efficacy |  | x 7 |  | x 7 |  | x 4 |
| Docetaxel | IC<sub>50</sub> (μM) | 93.6±8 | 32.6±7 | 158.6±5 | 92±13 | 178.2±21 | 144.8±16 |
|  | Efficacy |  | x 2.9 |  | x 1.7 |  | x 1.2 |

**Figure 3.**
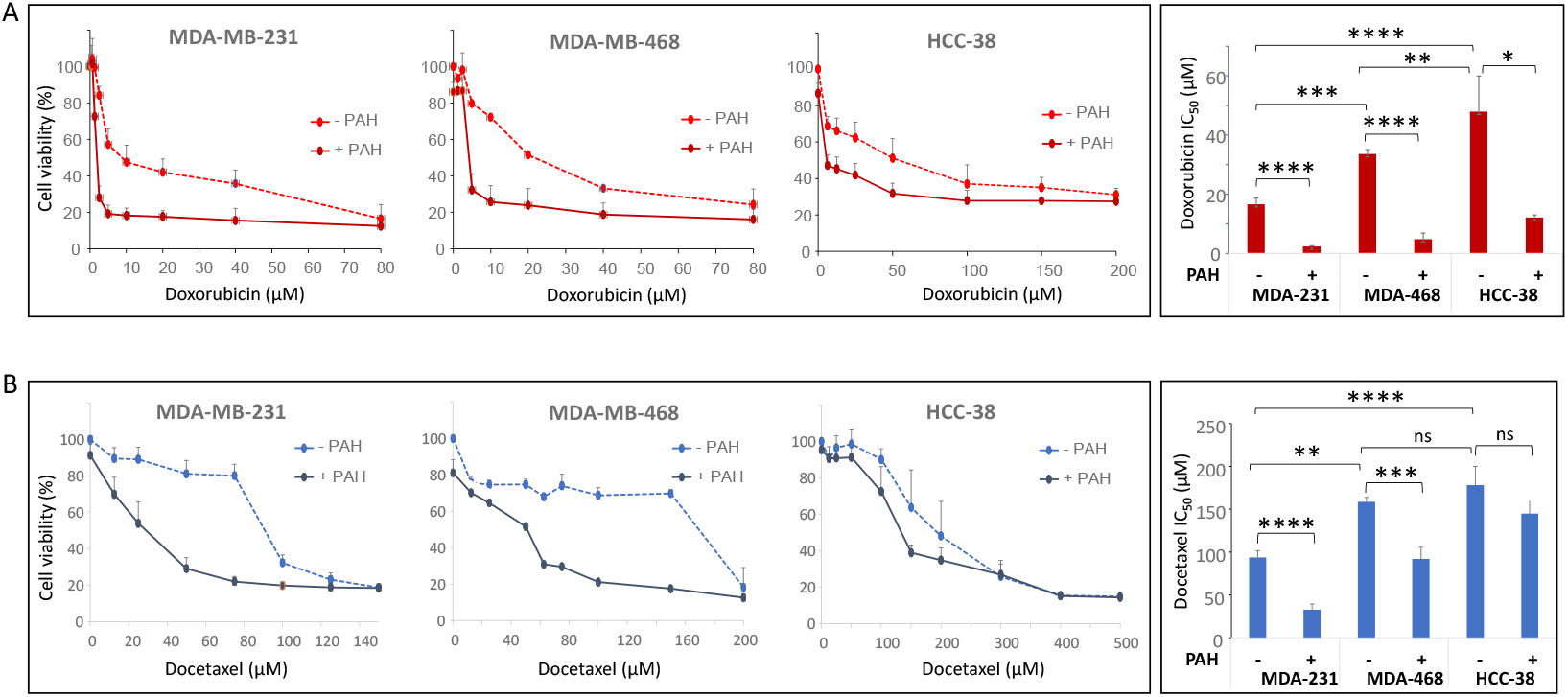
PTCH1 drug efflux inhibitor PAH increases the sensitivity of TNBC cells to chemotherapy. Cell viability was measured after 24 hours or 48 hours treatment with increasing concentration of docetaxel or doxorubicin respectively on MDA-MB-231, MDA-MB-468 and HCC-38 cell lines in the absence or the presence of 15µM PAH. IC_50_ values (corresponding to the concentration of chemotherapy inducing 50% of cell death) were calculated. Data reported are the mean ± SEM of at least 3 independent experiments. Significance is attained at P < 0.05 (*).

However, the effect of PAH is less pronounced on HCC-38 cells, while these cells express the highest amount of PTCH1. Interestingly, we found that HCC-38 cells expressed significantly higher amounts of P-glycoprotein (P-gp) and ABCG2 than MDA-MB-231 and MDA-MB-468 cells (Fig. 4). P-gp and ABCG2 are multidrug transporters from the ABC family well known to contribute to drug resistance of cancer cells. This could explain that PAH has less effect on HCC-38 cells than on MDA-MB-231 and MDA-MB-468.

**Figure 4.**
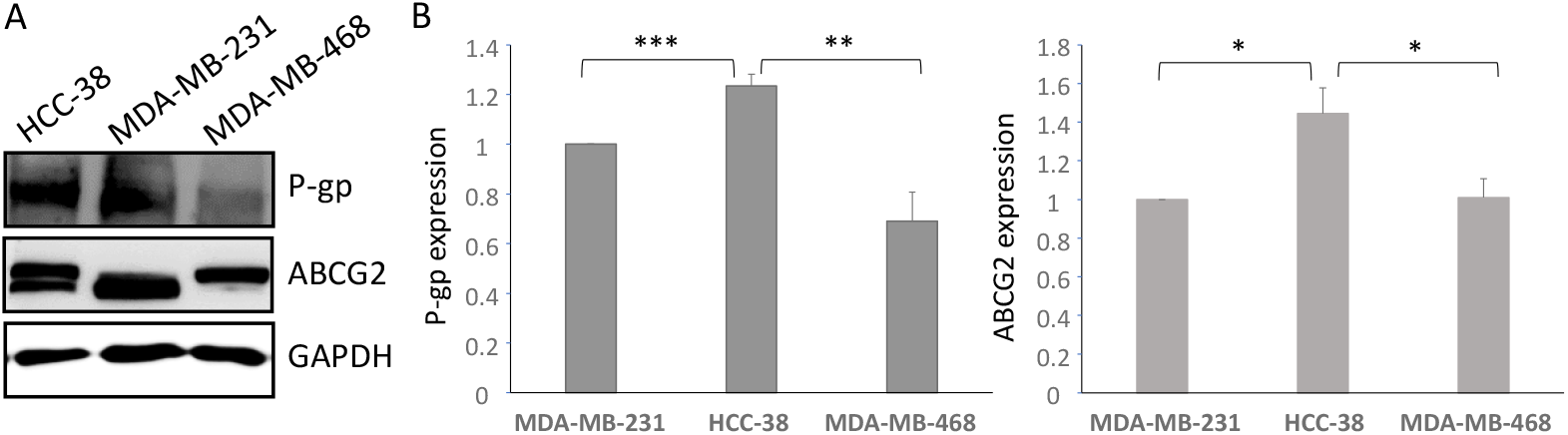
Other multidrug transporters are expressed in TNBC cell lines. **A**. P-gp and ABCG2 protein expression in three TNBC cell lines. Western-blot was performed on 50 µg extracts from each TNBC cell line (MDA-MB-231, HCC-38 and MDA-MB-468) with antibodies directed against P-gp or ABCG2 and GAPDH. **B**. P-gp, ABCG2 and GAPDH signals were quantified using ImageJ software. Data presented are the mean ± SEM of at least 3 independent experiments. Significance is calculated using Oneway ANOVA Turkey’s multiple comparisons test and attained at P < 0.05 (*). **P < 0.01; ***P < 0.001; ns P > 0.05.

We then tested the effect of PTCH1 drug efflux inhibition on MDA-MB-231 spheroids which more closely resemble to tumors than 2D cultures. We observed that the addition of 10 or 30 µM PAH destroyed spheroids at 250 and 200 µM docetaxel respectively, while these concentrations of docetaxel have no effect alone (Fig. 5). This experiment confirmed that PTCH1 drug efflux inhibition strongly increased the cytotoxicity of chemotherapy treatments on TNBC cells.

**Figure 5.**
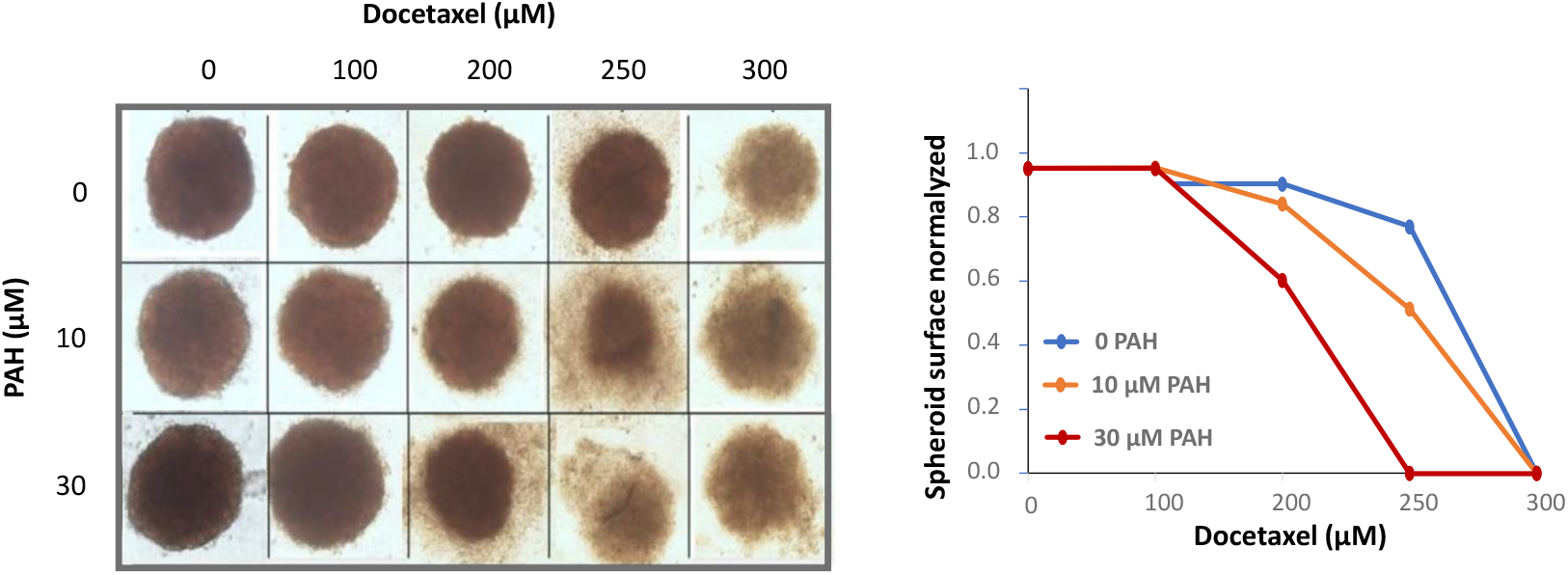
PAH increases docetaxel efficacy against TNBC spheroids. MDA-MB-231 cells were plated in ultra-low attachment surface 24 well plates in complete medium and treated with increasing concentrations of docetaxel in the presence of DMSO (control, 0 PAH), PAH 10 µM or 30 µM. After 2 weeks pictures were taken using Cytation 5 cell imaging system from Biotek. The surface of spheroids was calculated and reported after normalization on the condition 0 docetaxel for each concentration of PAH.

### PTCH1 drug efflux inhibition enhances the effect of chemotherapies on TNBC cells migration

In a previous study, we observed that PTCH1 is over-expressed in cells presenting cancer stem cell or persistent properties, and the present study highlighted that PTCH1 is expressed in circulating tumor cells from TNBC patients (Fig. 2). Cancer stem cells and CTC have a strong ability to migrate and form metastases. To know if the inhibition of PTCH1 drug efflux increased the cytotoxicity of chemotherapy against these cells, we performed migration assays. MDA-MB-231 and MDA-MB-468 cells were seeded on transwell membranes and treated with doxorubicin or docetaxel 24 or 48 hours respectively in the absence or the presence of PAH. The observation on a microscope of the cells that migrated to the bottom of the wells after staining with crystal violet revealed that the treatment with 1 µM doxorubicin or 20 µM docetaxel did not affect the number of cells in the wells in comparison to the non-treated wells (Fig. 6A), indicating that at these concentrations, doxorubicin and docetaxel had no effect on cell able to migrate. Remarkably, the addition of PAH at a concentration that has no effect alone strongly decreased the number of cells in the wells treated with 1 µM doxorubicin or 20 µM docetaxel, suggesting that the addition of PAH to the chemotherapies allowed to eliminate migrating cancer cells.

**Figure 6.**
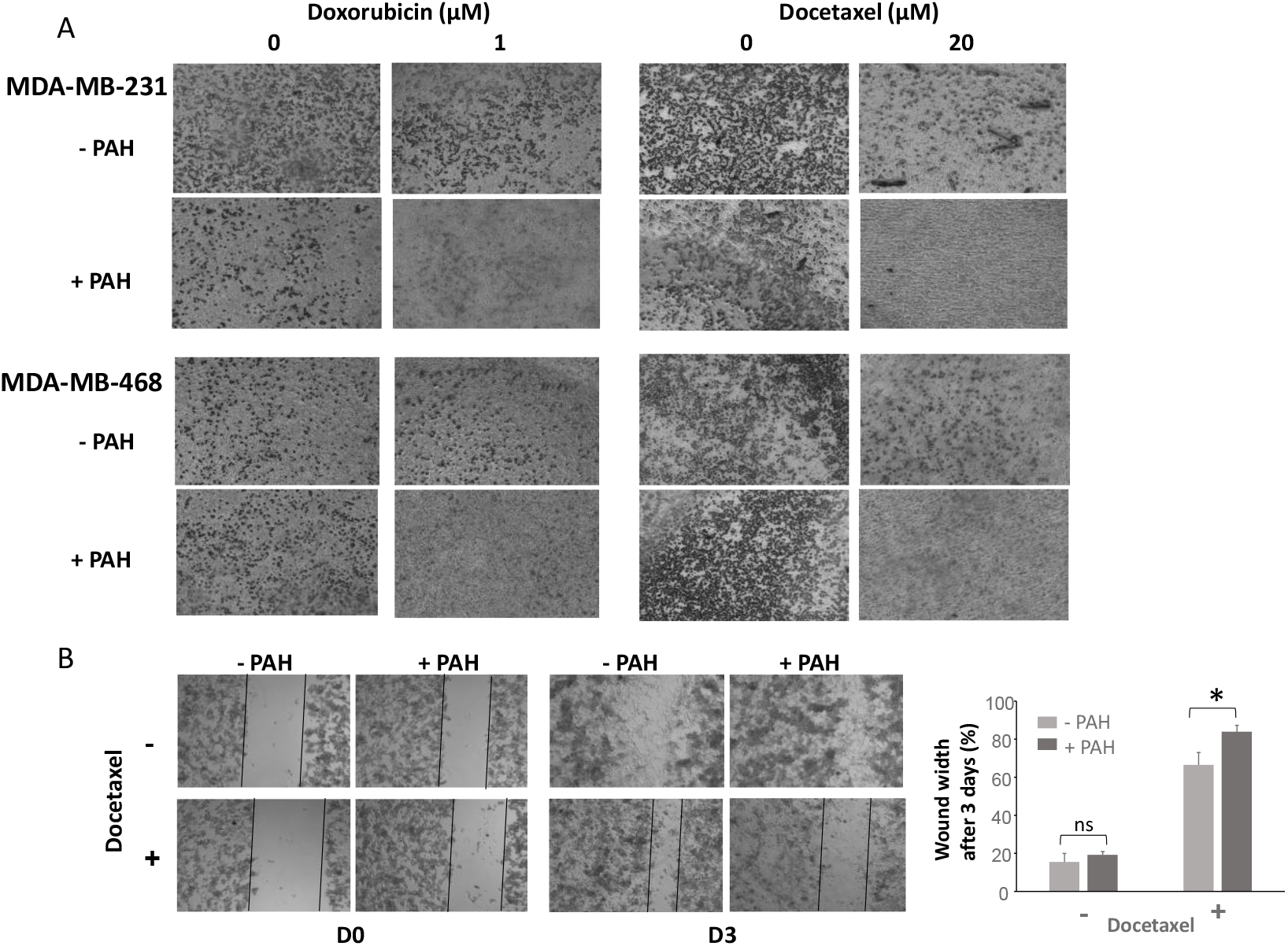
PAH addition to chemotherapy inhibits migration of TNBC cells. ***A***. 100000 cells were seeded on membrane from Transwell plates. Cells were treated 24 hours with doxorubicin +/-15µM PAH or 48 hours with docetaxel +/-15µM PAH. After fixation and staining with cristal violet, wells were observed on a microscope with X5 objective. ***B***. Migration was measured using a wound-healing assay. A wound was performed on MDA-MB-231 confluent cells seeded in 24 well plates. The medium was replaced by fresh medium containing 5 µM docetaxel in the absence of PAH, or the presence of 5 µM PAH. Two pictures were taken at two different points of each well immediately after wound, and 3 days after wound with 5X objective. The width of the wound was measured using ImageJ software and reported as final wound width/initial wound width in percentage. Data presented are the mean ± SEM of 3 independent experiments. Significance is attained at P < 0.05.

This conclusion was strengthened by wound-healing assays (Fig. 6B). Indeed, addition of 5 µM PAH to docetaxel treatment after a wound was performed on confluent MDA-MB-231 cells prevented more significantly the wound closure that docetaxel alone, confirming that PTCH1 inhibition allowed docetaxel to eliminate migrating cells. Note that 5 µM PAH has no effect by itself on the wound closure.

### PTCH1 drug efflux inhibition increases the concentration of doxorubicin in TNBC cells

When we treat cells with doxorubicin, which is naturally fluorescent, we observe a strong accumulation of doxorubicin in cells (Fig. 7A1). However, 30 min after removing doxorubicin from the medium, the fluorescence intensity in cells is strongly decreased indicating an efflux of doxorubicin out of cells (Fig. 7A2). The addition of PAH to the efflux medium after removing doxorubicin enabled relatively high fluorescence intensity to be maintained in the cells indicating that PAH inhibited the efflux of doxorubicin out of cells (Fig. 7A3). The efflux of doxorubicin after 30 min is around 50% in the absence of PAH, and significantly inhibited in the presence of PAH in the three TNBC cell lines (Fig. 7B). However, the effect of PAH is less pronounced on HCC-38. This is in good agreement with the results obtained in cell viability tests and confirms that other multidrug transporters could be involved in doxorubicin efflux from HCC-38 cells.

**Figure 7.**
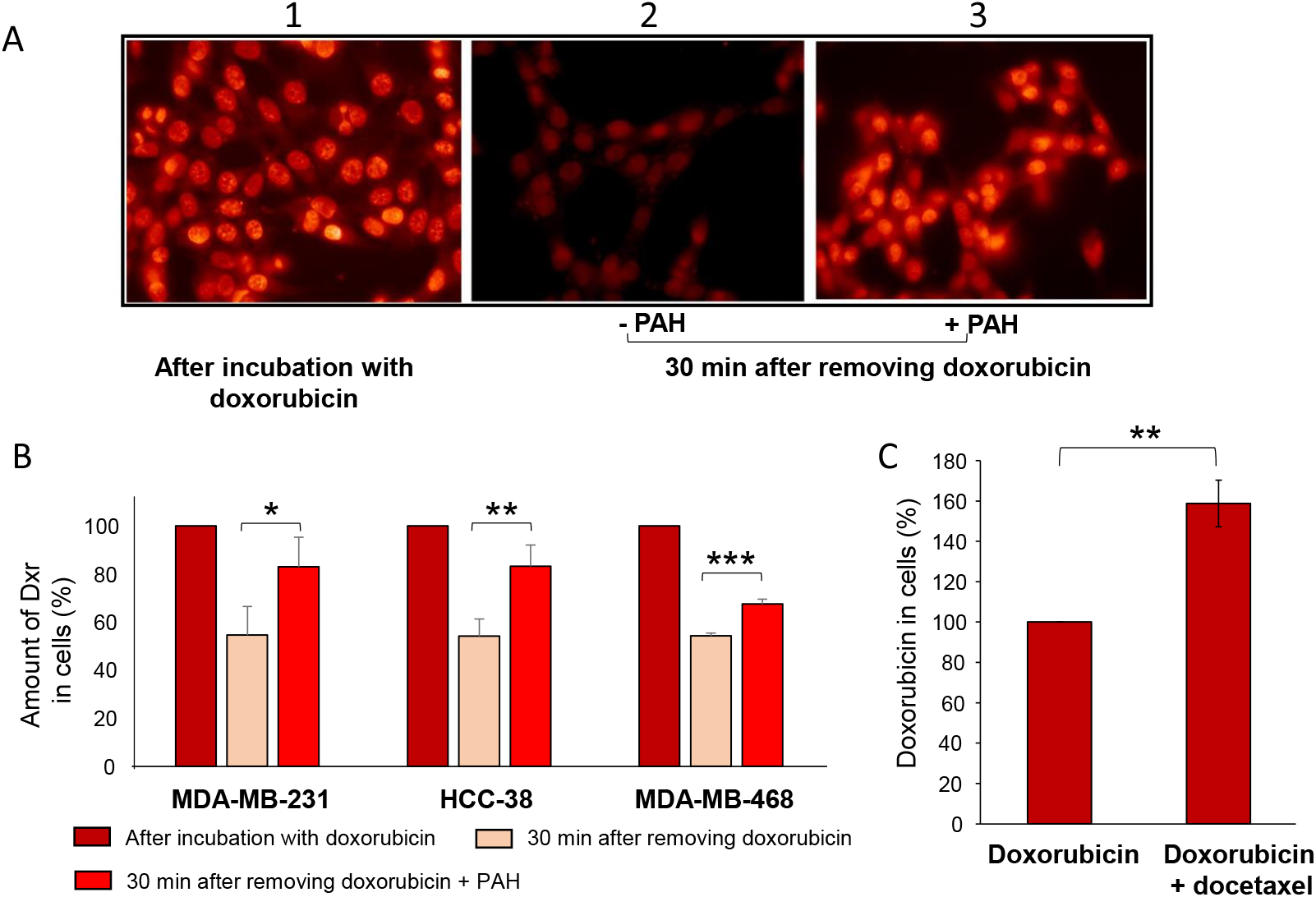
PAH inhibits doxorubicin efflux from TNBC cells. **A**. Cells on coverslip were incubated 2 hours with 10 µM doxorubicin, 3 coverslips were fixed, 6 coverslips were incubated with medium without doxorubicin +/- PAH during 30 min and fixed. Fluorescence of doxorubicin in cells was then observed by epifluorescence microscopy. **B**. the doxorubicin fluorescence in cells was quantified using ImageJ softwar. **C**. Docetaxel inhibits the accumulation of doxorubicin in MDA-MB-231 cells. Cells on coverslip were incubated with 10 µM doxorubicin or 10 µM doxorubicin and 50 µM docetaxel. Fluorescence of doxorubicin in cells was then observed by epifluorescence microscopy and quantified using ImageJ software. Histograms are the mean of at least 3 independent experiments. Significance is attained at P < 0.05 (*).

We also observed that the addition of an excess of docetaxel to doxorubicin increased doxorubicin accumulation in MDA-MB-231 cells (Fig. 7C), indicating that docetaxel inhibited the efflux of doxorubicin and that docetaxel is transported out of cells by PTCH1 like doxorubicin. This is in good agreement with the data showing that the expression of PTCH1 confers to yeasts a resistance to docetaxel (Sup Fig. 2) suggesting a role of PTCH1 in docetaxel efflux.

These results suggest that the increased cytotoxicity of doxorubicin and docetaxel observed in figures 3 and 5 could be due to the inhibition of their efflux from cancer cells by PTCH1.

## Discussion

The aim of this study was to evaluate the role of PTCH1 in the resistance to chemotherapy in breast cancer cells, and the effect of the inhibitor of its drug efflux activity, PAH, on the efficacy of chemotherapy treatments.

We first analyzed the expression of PTCH1 mRNA in cohorts of breast cancer patients, and observed that PTCH1 was expressed at the mRNAs level in all types of breast tumors, with more expression in TNBC (Fig. 1A,B). Our data revealed that PTCH1 mRNAs were also expressed in CTCs isolated from metastasis breast cancer patient blood and particularly in CTCs isolated from TNBC patients (Fig. 1C). Our analyses also highlighted that high Ptch1 mRNA expression is associated with poor prognosis (Fig. 1D, Sup Fig. 1). Our observations are in agreements with those reported in several studies (Jeng et al 2013, Wu et al 2022, Ozcan 2023). Indeed, using the Oncomine database, Wu and co-workers (2022) found that PTCH1 is expressed in breast cancer tissue and that PTCH1 expression level is associated with a poor prognosis in breast cancer patients. Moreover, analyses performed by Ozcan and co-workers (2023) on 5 GEO datasets and TCGA breast cancer data led them to conclude that PTCH1 is a key predictor of resistance to neoadjuvant chemotherapy in ER+/HER2-breast cancer.

TNBC is the most aggressive subtype of breast cancers. It is characterized by rapid disease progression, strong invasiveness, and high recurrence and metastasis rates, and is associated with a poor prognosis due to its limited treatment options. Despite recent advances in targeted therapy and immunotherapy, chemotherapies such as platinum agents (cisplatin, carboplatin), taxanes (paclitaxel and docetaxel), and anthracyclines (doxorubicin, epirubicin) remain the standard treatment option for TNBC across all stages (Lee 2023, Ou et al 2024). However, initial responses are mitigated with a high rate of chemoresistance and disease progression. A high percentage of TNBC patients experience relapse within 3–5 years following treatment (Garrido-Castro et al 2019, Li et al 2022). Compared to the other breast cancer subgroups, TNBC is enriched in a subpopulation of cells with self-renewal ability, termed breast cancer stem cells (BCSC) or tumor-initiating cells (TIC) that are drug-resistant and thought to be involved in the high relapse rate of TNBC patients (Park et al. 2019). Given that we previously showed that PTCH1 is over-expressed in a subpopulation of adrenocortical carcinoma cells displaying high chemotherapy resistance and TIC properties (Feliz Morel et al. 2022), and that PTCH1 is expressed in patient TNBC and in TNBC cell lines, we wondered whether PTCH1 was involved in the drug resistance in TNBC cells. In previous studies, we demonstrated that PTCH1 was able to efflux doxorubicin, and that PAH inhibited the doxorubicin efflux by PTCH1 in yeasts over-expressing PTCH1 and in melanoma cell lines expressing endogenous PTCH1 (Bidet et al 2012, Fiorini et al 2015, Signetti et al 2020). The inhibition of doxorubicin efflux from three TNBC cell lines endogenously expressing PTCH1 (MDA-MB-231, HCC-38 and MDA-MB-468) observed in the present work demonstrates that PAH is also able to inhibit doxorubicin efflux activity of PTCH1 in TNBC cell lines (Fig. 7). The fact that an excess of docetaxel inhibited doxorubicin efflux from a TNBC cell line suggested that docetaxel uses the same efflux pathway than doxorubicin (Fig. 7C).

In the present study, we showed that the expression of PTCH1 in yeasts conferred resistance to docetaxel, and that the addition of PAH to the culture medium inhibited this resistance (Sup Fig. 2B). These results demonstrate that PTCH1 is able to efflux docetaxel and to induce resistance to this chemotherapeutic drug. Our data revealed that PAH significantly enhanced the cytotoxic effect of doxorubicin and docetaxel on three TNBC cell lines grown in 2D (Fig. 3) and in 3D (spheroids) (Fig. 5), an *in vitro* model that more closely resemble to a tumor than adherent cells. Interestingly, the cell line showing the highest amount of PTCH1 (HCC-38) exhibited the strongest resistance to these two chemotherapies (Fig. 2B, Fig. 3, Table 1). However, PAH had a relatively low effect on HCC-38 in comparison to that induced on MDA-MB-231 and MDA-MB-468. We found that HCC-38 expressed higher amounts of P-glycoprotein (P-gp) and ABCG2 than MDA-MB-231 and MDA-MB-468 (Fig. 4). These two multidrug transporters from the ABC transporters family are known to trigger doxorubicin and docetaxel resistance (Mirzaei et al. 2022, Xu L et al 2025, Toyoda et al 2019). Despite its high expression in HCC-38, PTCH1 contributes probably less than P-gp and ABCG2 to chemotherapy resistance in HCC-38 than in MDA-MB-231 and MDA-MB-468. Moreover, we observed that the addition of PAH to docetaxel prevented TNBC cell migration (Fig. 6). PTCH1 being over-expressed in subpopulations of cancer stem cells with migration properties, the inhibition of its drug efflux activity by PAH made docetaxel toxic to these cells and allowed the elimination of migrating cells.

Taken together, our data suggest that PTCH1 is involved in the resistance of TNBC cells to chemotherapy, and that the use of an inhibitor of PTCH1 drug efflux during neoadjuvant or adjuvant therapy could prevent resistance of TNBC cells to treatment and metastases formation, and enhance the efficacy of the treatment against TNBC expressing PTCH1. Our study highlights the therapeutic potential of targeting PTCH1 using drug association strategies for the treatment of TNBC patients.

## Acknowledgments

We acknowledge the UCAGenomiX platform, partner of the National Infrastructure France Génomique, supported by the Commissariat Aux Grands Investissements (ANR-10-INBS-09-03, ANR-10-INBS-09-02).

## Author Contributions

Conception and design, I.M-V.; Development of methodology, S.C., S.K., T.B., R.W., E.D., A.T., S.A. and I.M-V.; Acquisition of data, S.C., S.K., T.B., R.W., E.D., A.T.; Analysis and interpretation of data, S.C., S.K., T.B., R.W., E.D., A.T., S.A. and I.M-V.; Writing – Original Draft, T.B., R.W. and I.M-V.; Writing –Review & Editing, T.B., R.W., M.F., F.L., S.A. and I.M-V..; Funding Acquisition, S.A. and I.M-V.; Resources, T.B., R.W., M.F., F.L., S.A. and I.M-V.; Supervision, I.M-V.

## Supplementary Figures

**Sup Figure 1.**
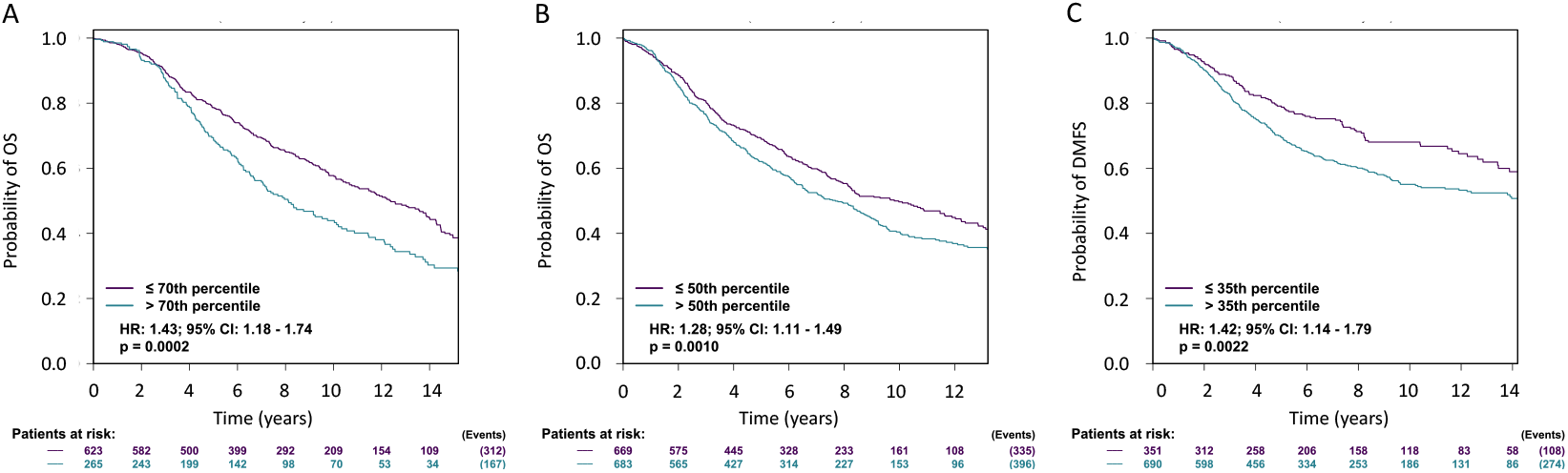
High PTCH1 mRNA expression is associated with a poorer prognosis in breast cancer. Distant Metastasis Free Survival (DMFS), disease-free survival (DFS) and overall survival (OS) data based on *PTCH1* mRNA expression were obtained from the intrinsic molecular subtypes’ prognostic analysis on ER+/HER2-high proliferative breast cancers performed on bc-GenExMiner v4.5 web portal and illustrated by Kaplan–Meier curves. The obtained Hazard Ratio (HR) with 95% confidence interval and log-rank P-values are shown.

**Sup Figure 2.**
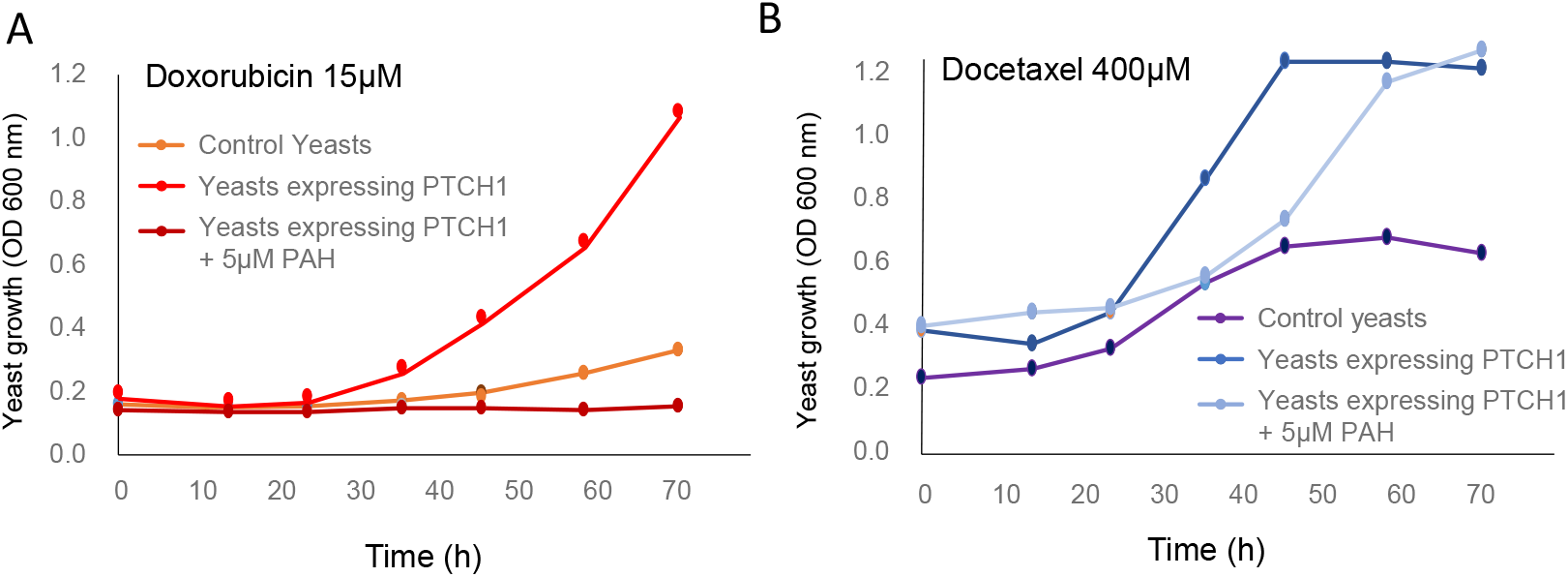
PTCH1 confers to yeasts a resistance to doxorubicin and docetaxel which is inhibited by PAH. Control yeasts and hPTCH1-expressing yeasts were grown in the presence of 15 µM doxorubicin (**A**) or 400µM docetaxel (**B**), supplemented or not with 5 µM of PAH. The growth of yeasts was measured by absorbance at 600 nm in function of time.

