## Supplementary figures 1 and 2 for "Inhibition of PTCH1 drug efflux activity enhances chemotherapy efficacy against triple negative breast cancer"

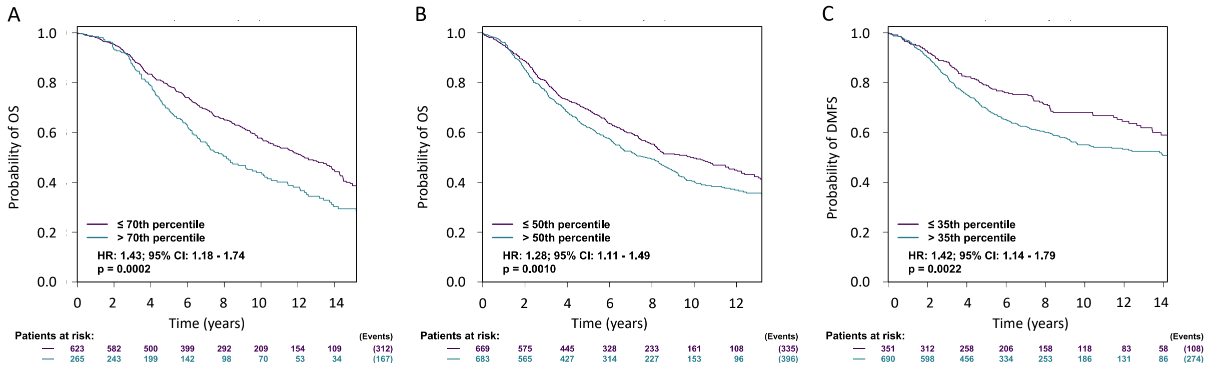


***Sup Figure 1. High PTCH1 mRNA expression is associated with a poorer prognosis in breast cancer.*** Distant Metastasis Free Survival (DMFS), disease-free survival (DFS) and overall survival (OS) data based on *PTCH1* mRNA expression were obtained from the intrinsic molecular subtypes' prognostic analysis on ER+/HER2- high proliferative breast cancers performed on bc-GenExMiner v4.5 web portal and illustrated by Kaplan–Meier curves. The obtained Hazard Ratio (HR) with 95% confidence interval and log-rank P-values are shown.

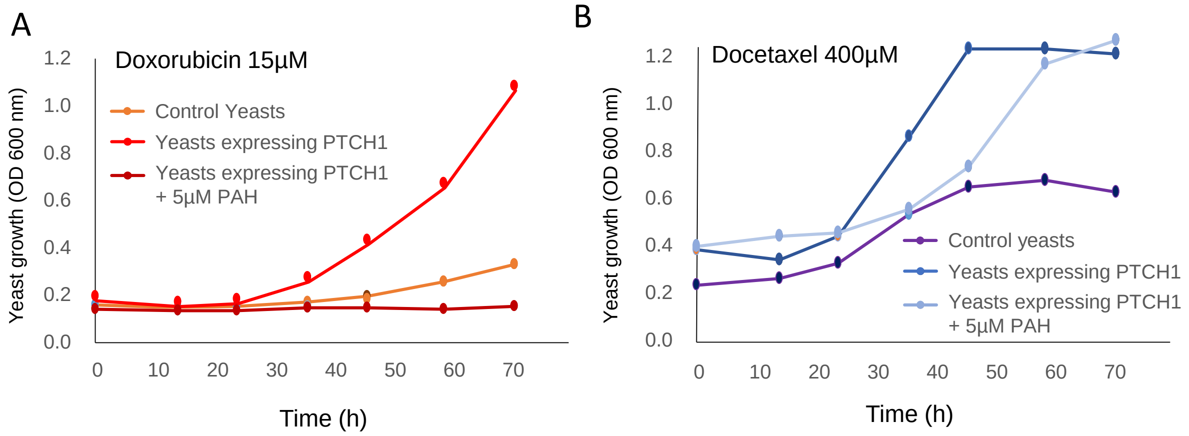


***Sup Figure 2. PTCH1 confers to yeasts a resistance to doxorubicin and docetaxel which is inhibited by PAH***. Control yeasts and hPTCH1-expressing yeasts were grown in the presence of 15 µM doxorubicin (**A**) or 400µM docetaxel (**B**), supplemented or not with 5 µM of PAH. The growth of yeasts was measured by absorbance at 600 nm in function of time.
